# A gut commensal protist protects against virus-mediated loss of oral tolerance

**DOI:** 10.1101/2022.06.21.497012

**Authors:** Magdalena Siller, Yanlin Zeng, Luzmariel Medina Sanchez, Pamela H. Brigleb, Kishan A. Sangani, Mohit Rana, Lauren Van Der Kraak, Surya P. Pandey, Mackenzie J. Bender, Britney Fitzgerald, Lee Hedden, Kay Fiske, Gwen M. Taylor, Syed A Rahman, Heather J. Galipeau, Steven J. Mullet, Stacy G. Wendell, Simon C. Watkins, Premysl Bercik, Jishnu Das, Marlies Meisel, Bana Jabri, Terence S. Dermody, Elena F. Verdú, Reinhard Hinterleitner

## Abstract

Loss of oral tolerance (LOT) to gluten, characterized by a T helper 1 (Th1) gluten-specific immune response, is a hallmark of celiac disease (CeD) and can be triggered by enteric viral infections. We hypothesized that certain gut microbes have the capacity to protect against virus-mediated LOT. By using our previously defined reovirus-mediated LOT CeD model, we discovered that the gut colonizing protist *Tritrichomonas* (T.) *arnold* promotes oral tolerance and protects against reovirus-mediated LOT by suppressing the reovirus-induced proinflammatory program of dietary-antigen-presenting CD103^+^ dendritic cells. Importantly, *T. arnold* did not affect antiviral host immunity, suggesting that *T. arnold*-mediated protection against T1L-induced LOT is not attributable to differences in antiviral host responses. Additionally, using gnotobiotic mice, we found that *Tritrichomonas arnold* colonization is sufficient to protect against reovirus-mediated LOT in the absence of the microbiota. Mechanistically, we show that *Tritrichomonas arnold* colonization restrains reovirus-induced inflammatory responses in dendritic cells and thus limit their ability to promote Th1 immune responses *ex vivo*. Finally, our studies using human stool samples support a role for *Tritrichomonas* sp. colonization in protecting against development of CeD. This study will motivate the design of effective therapies to prevent LOT to gluten in at-risk individuals and to reinstate tolerance to gluten in CeD patients.

**One Sentence Summary:** *Tritrichomonas arnold* protects against virus-mediated loss of oral tolerance to gluten and is underrepresented in celiac disease patients.

## Main Text

Celiac disease (CeD) is an immune disorder in which genetically susceptible individuals expressing the human leukocyte antigen (HLA) DQ2 or DQ8 molecules display an inflammatory T helper 1 (Th1) immune response against dietary gluten present in wheat, barley, and rye that results in loss of oral tolerance to gluten (LOT) (*1–3*). The HLA-DQ2-or HLA-DQ8-restricted Th1 response against gluten, a hallmark of CeD, initiates CeD pathogenesis, which is characterized by cytotoxic intraepithelial CD8^+^ lymphocyte-mediated tissue destruction resulting in villous atrophy (*4, 5*). Intriguingly, although approximately 30% of the human population carry HLA DQ2 or DQ8, only 1% develops CeD (*1*), suggesting that additional environmental factors contribute to CeD pathogenesis. Viruses have been implicated as a potential environmental trigger in CeD pathogenesis (*6–9*). Concordantly, we discovered that reovirus, a largely avirulent pathogen that elicits protective immunity, can nonetheless promote LOT by suppressing regulatory T cell (Treg) conversion and promoting inflammatory Th1 immunity to dietary antigen. Mechanistically, the human reovirus isolate, type 1 Lang (T1L) mediates LOT by endowing dietary antigen-presenting dendritic cells (DCs) with proinflammatory properties in the mesenteric lymph nodes (mLN), the inductive site of oral tolerance (*10*).

In the course of our studies of reovirus T1L-mediated inflammatory responses to dietary antigen (*10*), we reproduced our findings using C57BL/6 mice commercially acquired from Jackson Laboratories (JAX). However, we did not to observe T1L-mediated inflammatory responses to dietary antigen in aged-and sex-matched C57BL/6 mice bred in our vivarium (“in-housed” mice originally acquired from JAX) using an oral tolerance assay (Fig. 1A-C **and** fig. S1A-D). T1L-mediated CD4^+^ Th1 immune responses specific for dietary antigen, characterized by elevated expression of IFNγ and Tbet, were absent in in-housed mice compared with JAX mice (Fig. 1A-C **and** fig. S1B-D). Untreated and T1L-infected in-housed mice displayed a significant expansion of Treg compared with untreated and T1L-infected JAX mice, respectively (Fig. 1A-C **and** fig. S1B-D), suggesting that environmental factors present in in-housed but not JAX mice mediate protection against T1L-mediated LOT. Intriguingly, T1L infection promoted host Tbet^+^ CD4^+^ T cell responses that were comparable between in-housed mice and JAX mice and concordant with our previous findings ((fig. S1E-G) and (*10*)), indicating that host Th1 cell responses to T1L infection are uncoupled from dietary-antigen-specific Th1 cell responses in in-housed mice. Dietary antigen uptake takes place in the small intestine (*11, 12*). Examination of the small intestine revealed a significantly enlarged small intestine and a substantial expansion of tuft and goblet cells in in-housed mice relative to JAX mice, independent of T1L infection (Fig. 1D-F). These findings suggest an elevated type-2 immune response, which is usually observed in mice infected with helminths (*13, 14*) or colonized with the gut commensal protest *Tritrichomonas* (*15*). As helminth infections are unlikely based on our specific-pathogen free (SPF) mouse facility status, we examined animals for *Tritrichomonas* in cecal contents. This analysis revealed the presence of *Tritrichomonas* in in-housed mice but not mice acquired from JAX (Fig. 1G). Internal transcribed spacer (ITS) sequencing identified this protist as an uncharacterized *Tritrichomonas* sp. isolate with the accession number MF375342 and with 86%, 84%, and 85% homology to the ITS of *Tritrichomonas muris*, *Tritrichomonas musculis*, and *Tritrichomonas rainier*, respectively ((fig S2A-B) and (*15–17*)). Although the precise taxonomic relationship of these tritrichomonads remains to be determined, for clarity we hereafter refer to the University of Pittsburgh isolate as *T. arnold*.

**Fig. 1.**
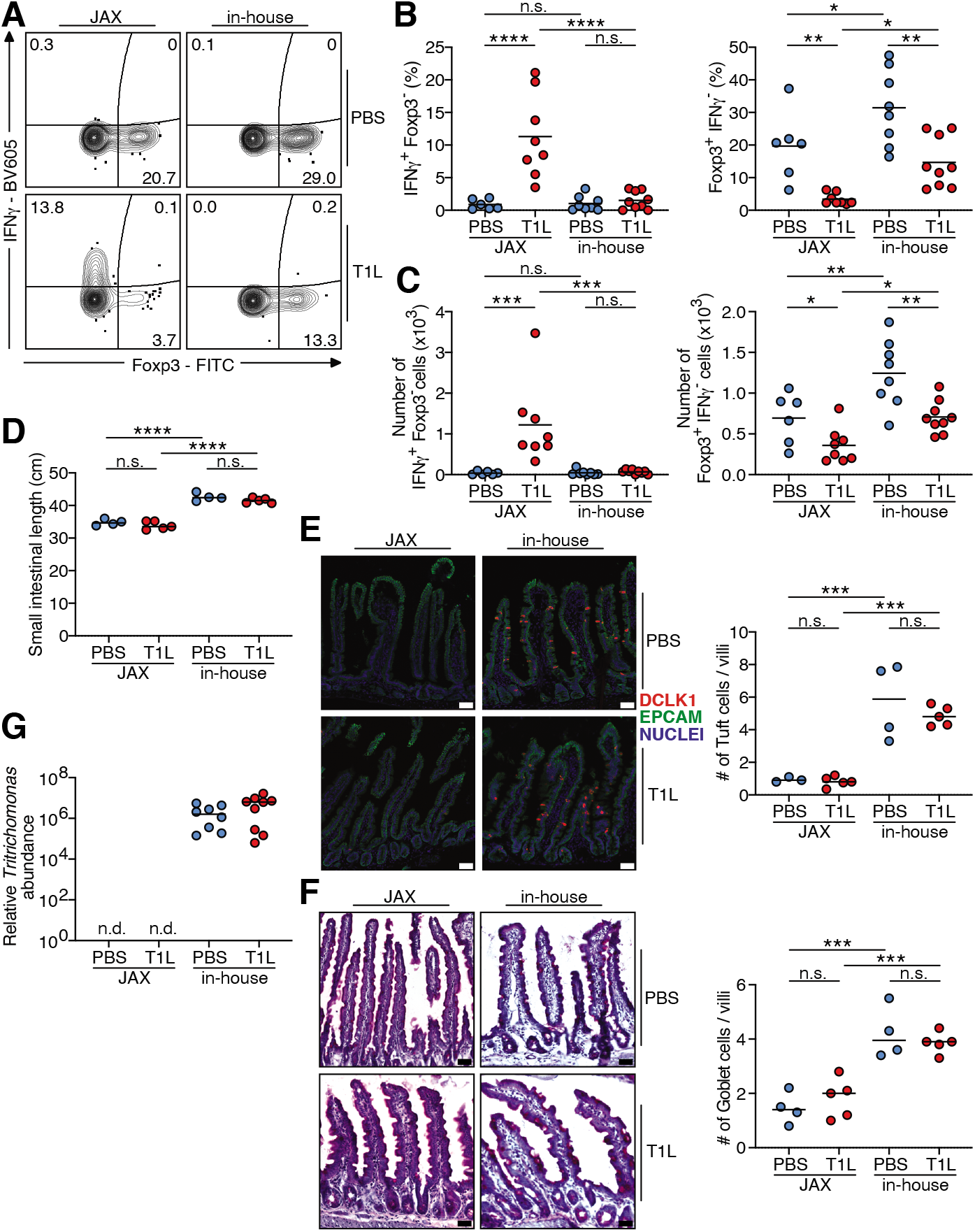
Suppression of T1L-induced loss of oral tolerance is associated with the presence of *Tritrichomonas* in in-housed mice. **(A-C)** Naïve OT-II^+^ CD45.1^+^ CD4^+^ T cells were transferred into WT CD45.2^+^ mice. One day after transfer mice were inoculated perorally with 10^9^ PFU of T1L or PBS and fed an OVA-containing diet for 6 days. The intracellular expression of Foxp3 and IFNγ in transferred OT-II^+^ CD45.1^+^ CD4^+^ T cells in the mLN was evaluated by means of flow cytometry. (A) Representative contour plots, (B), percentages and (C), absolute numbers are shown. **(D)** Small intestine length **(E)** Presence of tuft cells in the jejunum visualized by the expression of DCLK1 in red; epithelial cells (EPCAM) in green; DAPI in blue. **(F)** Total number of PAS^+^ goblet cells per villi in the jejunum was determined. (E-F) Representative images (left panel), quantification (right panel); Scale bars, 50 µm. **(G)** *Trichomonas* colonization was quantified by qPCR using DNA isolated from cecal contents. 28S normalized to the host murine *Ifnb1* gene. (B-G) Data are representative of two independent experiments (individual dots represent n-numbers). Center is mean, one-way ANOVA, Tukey’s post hoc test. \**P* < 0.05, \*\**P* < 0.01, \*\*\**P* < 0.001, \*\*\*\**P* < 0.0001, n.s. not significant.

In experiments to better understand how *T. arnold* colonization affects mucosal CD4^+^ T cell responses to dietary antigens in the absence of enteric T1L infection (fig. S3A), we found that *T. arnold* colonization of JAX WT mice promoted significant enhancement of dietary-antigen-specific Tregs relative to controls (fig. S3B-D). Although *T. arnold* induced mucosal type-2 responses (Fig. 1E-F), we did not observe *T. arnold* -induced Th2 responses against dietary antigen (fig. S3E). Of note, *T. arnold* colonization neither promoted Th1 nor Th17 cell differentiation against dietary antigen (fig. S3B-D **and** S3F). These findings suggest that *T. arnold* selectively drives dietary-antigen-specific Treg differentiation.

To test the sufficiency of *T. arnold* in protecting against T1L-mediated inflammatory responses to dietary antigen, we orally inoculated JAX WT mice with *T. arnold*. Twelve days post *T. arnold* colonization, we conducted an oral tolerance assay (fig. S4A). Oral gavage of *T. arnold* into 6-week-old JAX WT mice resulted in colonization levels comparable to naturally colonized in-housed mice (fig. S4B). Confirming our previous observations (*10*), T1L blocked Treg responses and promoted Th1 immune responses specific for dietary antigen in PBS-treated mice (Fig. 2A-C). Strikingly, *T. arnold* efficiently counteracted both T1L-mediated Treg suppression and Th1 immune responses against dietary antigen (Fig. 2A-C). Importantly, *T. arnold* did not affect viral replication, anti-reovirus antibody titers, and host type-1 interferon responses (Fig. 2D-F), suggesting that *T. arnold*-mediated protection against T1L-induced LOT is not attributable to differences in antiviral host responses. These findings indicate that *T. arnold* colonization promotes tolerogenic responses to dietary antigen that can suppress virus-mediated LOT.

**Fig. 2.**
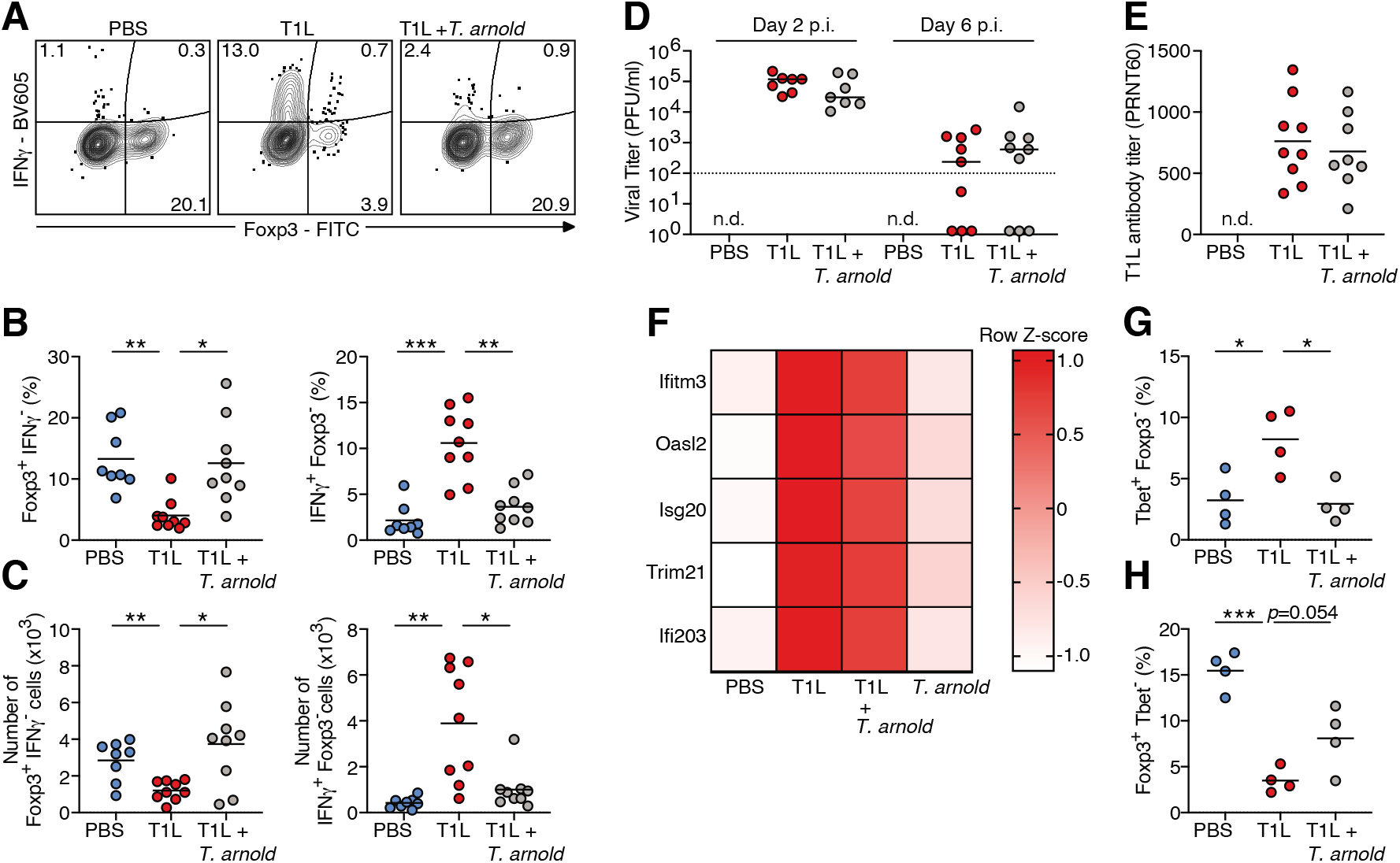
*T. arnold* protects against T1L-mediated loss of oral tolerance without impacting antiviral immunity and independent of the microbiota. **(A-F)** WT CD45.2^+^ mice were inoculated perorally with 10^6^ *T. arnold* or PBS for 12 days prior to T cell transfer and/or T1L infection. (A-C) Naïve OT-II^+^ CD45.1^+^ CD4^+^ T cells were transferred into WT CD45.2^+^ mice. One day after transfer mice were inoculated perorally with 10^9^ PFU of T1L or PBS and fed an OVA-containing diet for 6 days. The intracellular expression of Foxp3 and IFNγ in transferred OT-II^+^ CD45.1^+^ CD4^+^ T cells in the mLN was evaluated by means of flow cytometry. (A) Representative contour plots, (B) percentages and (C), absolute numbers are shown. **(D)** 48 hours and 6 days post T1L infection, a 3cm section of ileum was resected and viral titers were determined by plaque assay. Plaque-forming unit (PFU). Dotted line indicates detection limit. **(E)** At 18 days post T1L infection sera were collected, heat inactivated, and used for a plaque-reduction neutralization assay. (D-E) Center is median. **(F)** RNA-sequencing of mLN 48 hours post T1L infection. Heat map of selected type-1 interferon inducible genes (ISGs). The scale represents the Row Z-score. (n = 4 all groups). **(G-H)** Germ-free CD45.2^+^ WT mice were inoculated perorally with 10^6^ *T. arnold* or PBS for 12 days prior to OT-II T cell transfer. One day after transfer mice were inoculated perorally with 10^9^ PFU of T1L or PBS and fed 1.5% OVA in drinking water for 6 days. The intracellular expression of (G) Tbet and (H) Foxp3 in transferred OT-II^+^ CD45.1^+^ CD4^+^ T cells in the mLN was evaluated by means of flow cytometry. Percentages are shown. (B-E, G-H) Data are representative of two independent experiments (individual dots represent n-numbers). (B-C, G-H) Center is mean, one-way ANOVA, Tukey’s post hoc test. \**P* < 0.05, \*\**P* < 0.01, \*\*\**P* < 0.001.

We next sought to determine how *T. arnold* confers the protective effect against T1L-mediated LOT. One possibility is that *T. arnold* influences oral tolerance indirectly by modulation of the microbiota. The short chain fatty acid (SCFA) butyrate produced by specific bacteria promotes tolerogenic immune responses and Treg differentiation (*18*). *Tritrichomonas* does not produce butyrate (*17*). Of note, *T. arnold* colonization did not alter concentrations of the SFCA butyrate, acetate, and propionate measured in cecum contents, suggesting that *T. arnold*-mediated protection against virus-induced LOT is independent of SCFA production by gut bacteria (fig. S5A-C). We directly tested whether the microbiota is required for the protective effects of *T. arnold* on T1L-induced LOT. T cell conversion assays conducted using germ-free mice showed that *T. arnold* protected from T1L-mediated LOT (Fig 2G-H), implying that *T. arnold* is sufficient to protect from T1L-mediated LOT independent of the microbiota.

Similar to studies using other *Tritrichomonas* sp. (*15, 17*), *T. arnold* colonization of JAX mice for 2 weeks resulted in significantly elevated type-2 immunity in the small intestine, characterized by tuft and goblet cell hyperplasia (fig. S6A-B). *Tritrichomonas* sp. promotes type-2 immunity by releasing succinate, a carboxylic acid and fermentation product of *Tritrichomonas* sp. from ingestion of complex polysaccharides. Succinate is sensed by succinate-receptor-expressing tuft cells and is sufficient to promote IL-25-mediated tuft and goblet cell expansion and ILC2 activation (*17, 19, 20*). As observed previously (*17, 19, 20*), succinate supplementation in drinking water was sufficient to induce type-2 immune responses in the small intestine comparable to *T. arnold* colonization in *Tritrichomonas*-free mice (fig. S6A-D). Next, we tested whether succinate supplementation in drinking water is sufficient to promote oral tolerance and protect against T1L-mediated inflammatory responses to dietary antigen. Mice were administered succinate in drinking water for 12 days prior to an OT-II conversion assay (fig. S6E). In contrast to *T. arnold*, succinate alone was neither sufficient to promote Treg responses to dietary antigen nor capable of protecting against T1L-mediated inflammatory responses to dietary antigen (fig. S6F-G). Of note, succinate failed to influence viral replication (fig. S6H). These data indicate that *T. arnold*-mediated oral tolerance is independent of the capacity of the protist to release succinate. Goblet cells have been implicated in dietary antigen uptake and oral tolerance (*21, 22*). Although we observe an expansion of goblet cells in *T. arnold*-colonized mice (fig. S6B), the fact that succinate supplementation induces similar levels of goblet cell expansion (fig. S6D) suggests that goblet cell expansion alone cannot explain the immunologic tolerance to dietary antigen induced by *T. arnold*.

*T. musculis* promotes IL-18 release by colonocytes, which confers protection against *Salmonella typhimurium* infection (*16*). To determine whether IL-18 functions in the protective effects of *T. arnold* in reovirus-induced LOT, we conducted an OT-II conversion assay using IL-18 deficient mice. *T. arnold* protected against T1L-induced LOT to a comparable extent in IL-18 deficient mice and WT controls, suggesting that IL-18 is dispensable for the effects of *T. arnold* on dietary-antigen-specific immune responses (fig. S7A-B). Collectively, we show that *T. arnold* colonization promotes oral tolerance and prevents T1L-mediated LOT independent of antiviral host responses, succinate sensing, IL-18 signaling, and the microbiota.

DCs orchestrate protective or tolerogenic T cell responses to reovirus infection and dietary antigen, respectively (*23–25*). In Peyer’s patches (PP), protective immunity to T1L is initiated by CD103^-^ CD11b^+^ DCs processing viral antigen (*23*). In contrast, in mLN, tolerogenic CD103^+^ DCs present dietary antigen and induce oral tolerance (*11*). In the mLN, dietary-antigen-presenting CD103^+^ CD11b^-^ CD8α^+^ DCs have the highest tolerogenic potential (*10, 24, 26*) but also drive Th1 responses to enteric pathogens (*10, 25, 27, 28*). T1L infection induces an IRF1-mediated proinflammatory program in CD103^+^ CD11b^-^ CD8α^+^ DCs, characterized by IL-12 production, activation of the costimulatory molecule CD86, and increased *Il27* gene expression, thereby promoting LOT (*10*). Hence, based on our finding that *T. arnold* blocks T1L-mediated inflammatory responses to dietary antigen, we determined whether *T. arnold* modulates protective or tolerogenic DC responses in PP and mLN, respectively. Analysis of DCs in PP and mLN revealed that T1L infection induces inflammatory DC activation as evidenced by IL-12 production and upregulation of CD86 at both sites (Fig. 3A-F **and** fig. S8A-B). Strikingly, *T. arnold* colonization specifically suppressed T1L-mediated IL-12 production and expression of CD86 and *Il27* in mLN DCs without substantially affecting T1L-mediated DC activation in PP (Fig. 3A-G **and** fig. S8A-B). CD103^+^ DCs in the small intestinal *lamina propria* take up dietary antigen and migrate to mLN to present dietary antigen to CD4^+^ T cells (*24, 26*). *T. arnold* colonization did not affect the overall composition of mLN DC subsets, efficiency of dietary-antigen uptake by mLN DCs or OT-II T cell proliferation (fig. S8C-E). Thus, the amount of dietary antigen available at inductive sites of oral tolerance is not influenced by the presence of *T. arnold*. Therefore, specific suppression of the T1L-mediated inflammatory program in dietary-antigen-presenting DCs by *T. arnold* is consistent with our findings that *T. arnold* prevents T1L-mediated LOT without affecting the antiviral host response.

**Fig. 3.**
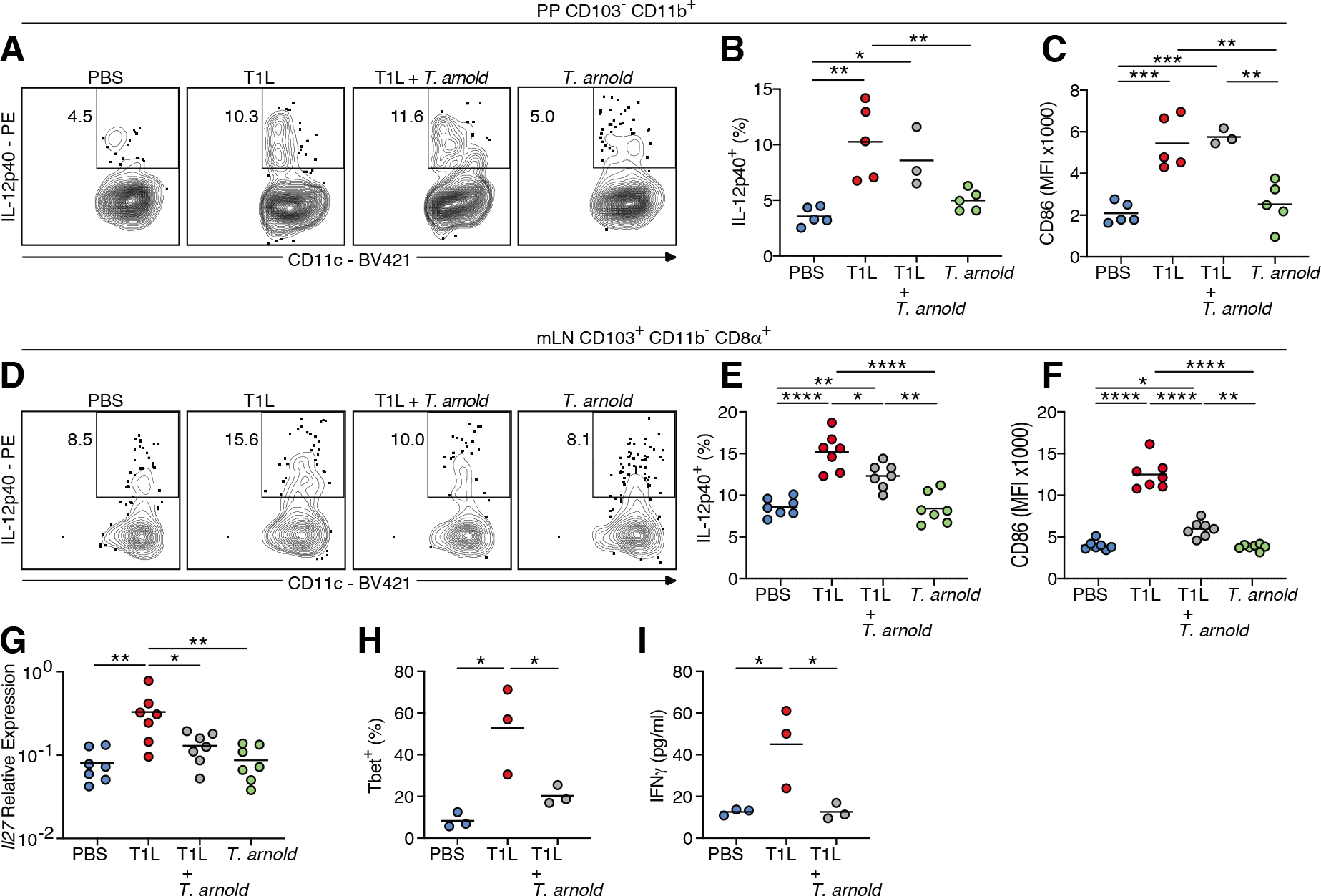
*T. arnold* suppresses the T1L-mediated proinflammatory program in dietary-antigen-presenting dendritic cells. **(A-I)** WT mice were inoculated perorally with 10^6^ *T. arnold* or PBS for 12 days followed by peroral inoculation with 10^9^ PFU of T1L or PBS for 48 hours. (A-C) The expression of (A-B) IL-12p40 and (C) CD86 on gated MHC-II^+^ CD11c^+^ CD103^-^ CD11b^+^ PP DCs was evaluated by means of flow cytometry. (D-F) The expression of (D-E) IL-12p40 and (F) CD86 on gated MHC-II^+^ CD11c^+^ CD103^+^ CD11b^-^ CD8α^+^ mLN DCs was evaluated by means of flow cytometry. (A-F) Representative dot plots, percentages, and MFI are shown. (G) *Il27* gene expression in the mLN was analyzed by means of RT-PCR. (H-I) Purified mLN DCs were co-cultured with naïve OT-II CD4^+^ T cells for 3 days in the presence of OVA peptide. (H) Intracellular expression of Tbet in OT-II CD4^+^ T cells was evaluated by means of flow cytometry. (I) Levels of IFNγ in the co-culture supernatants were evaluated by means of electrochemiluminescence. (A-I) Data are representative of two independent experiments (individual dots represent n-numbers). Center is mean, one-way ANOVA, Tukey’s post hoc test. \**P* < 0.05, \*\**P* < 0.01, \*\*\**P* < 0.001, \*\*\*\**P* < 0.0001.

The capacity of *T. arnold* to suppress T1L-mediated inflammatory DC responses in mLN encouraged us to determine whether DCs derived from mLN of *T. arnold*-colonized mice are sufficient to suppress T1L-induced inflammatory Th1 cell responses *ex vivo*. Remarkably, co-cultures of mLN-derived DCs isolated 48 hours after T1L inoculation in the presence or absence of *T. arnold* colonization with naïve CD4^+^ OT-II T cells and OVA peptide revealed that the presence of *T. arnold* suppressed the T1L-mediated Th1 response by modulating DCs responses (Fig. 3H-I **and** fig S8F). Together, these findings demonstrate that *T. arnold* colonization is sufficient to suppress a T1L-mediated inflammatory program in dietary-antigen-presenting DCs.

To determine the relevance of these findings to CeD, we analyzed the effect of *T. arnold* colonization on T1L-mediated LOT to gluten in transgenic mice expressing the CeD-predisposing HLA molecule DQ8 (DQ8tg mice) (*10*). Whereas T1L induced LOT to gluten in mice without *T. arnold* colonization, the presence of *T. arnold* prevented T1L-mediated LOT to gluten in DQ8tg mice, as assessed by the presence of anti-gliadin IgG2c antibodies and the development of a Th1 delayed-type hypersensitivity reaction (Fig. 4A-B **and** fig. S9A). In addition to intestinal environmental conditions favoring Th1 immunity against dietary antigen, TG2 activation is thought to promote CeD pathogenesis by increasing the affinity of gluten peptides for HLA-DQ2 and HLA-DQ8 molecules through posttranslational modifications (*1–3*). T1L infection induces TG2 activation (*10*). To test whether *T. arnold* suppresses T1L-mediated TG2 activation, we quantified incorporation of 5-(biotinamido)-pentylamine (5BP), a small-molecule TG2 activity probe. We found that *T. arnold* effectively suppressed T1L-mediated TG2 activation (Fig. 4C), suggesting that *T. arnold* prevents LOT and also prevents the modification of gluten peptides. Thus, in a CeD-relevant mouse model, *T. arnold* prevents T1L-mediated abrogation of oral tolerance to gluten and T1L-mediated TG2 activation, suggesting that *T. arnold-*related strains may protect against LOT to gluten and development of CeD.

**Fig. 4.**
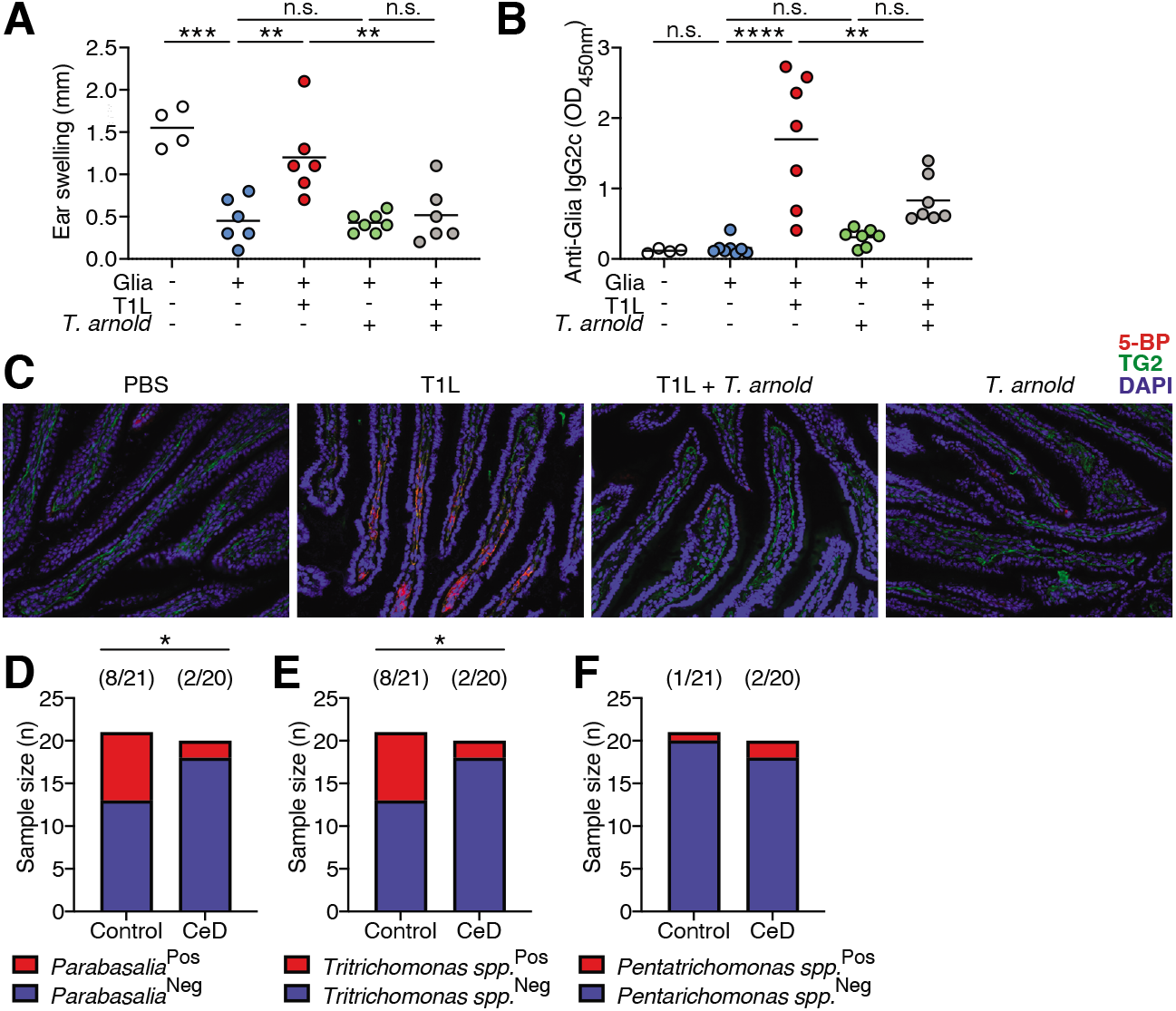
*Parabasalia* protect against loss of oral tolerance to gluten and are underrepresented in celiac disease patients compared to healthy controls. (**A** and **B**) DQ8tg mice were inoculated perorally with 10^6^ *T. arnold* or PBS for 12 days followed by peroral inoculation with 10^9^ PFU of T1L or PBS at the initiation of an oral tolerance/delayed type hypersensitivity protocol. Mice were fed orally with gliadin (Glia) or PBS for 2 days and then immunized subcutaneously with a CFA-Glia emulsion. (A) Levels of Glia-specific IgG2c antibodies in the serum were quantified at day 27 by means of enzyme-linked immunosorbent assay (ELISA). (B) On day 29 and 39, mice were challenged subcutaneously with Glia, and the degree of ear swelling was determined 48 hours after the second challenge. Data are representative of two independent experiments (n = 4-7 per group). Centre is mean, one-way ANOVA, Tukey’s post hoc test. **(C)** DQ8tg mice were colonized with 10^6^ *T. arnold* for 12 days prior to T1L infection. DQ8tg mice were inoculated perorally with 10^9^ PFU of T1L or PBS. Mice were euthanized at 18 hours after infection and jejunum was collected and frozen in optimal cutting temperature compound. (C) Representative images from stained frozen sections of the jejunum are shown. Scale bars, 100µm. Staining with 4’,6-diamidino-2-phenylindole (DAPI) is shown in blue, TG2 protein is shown in green, and TG2 enzymatic activity as assessed by means of 5BP cross-linking is shown in red. **(D-F)** DNA was isolated from stool samples collected from healthy volunteers and active CeD patients and subjected to ITS PCR-DNA sequence analysis. Frequency of (D) *Parabasalia* (E) *Tritrichomonas* sp. and (F) *Pentatrichomonas hominis* detected in healthy volunteers and active CeD patients (detected (Pos), not detected (Neg)). Two-sided Chi-square test. \**P* < 0.05, \*\**P* < 0.01, \*\*\**P* < 0.001, \*\*\*\**P* < 0.0001.

Based on our findings about the protective role of *T. arnold* in a CeD mouse model, we hypothesized that protists of the *Parabasalia* family, including *Tritrichomonas* and other human gut colonizing protists such as *Dientamoeba fragilis* and *Pentatrichomonas hominis* (*16*), protect against LOT and CeD development in humans. To investigate a role for *Parabasalia* in CeD, we isolated DNA from stool samples of healthy individuals and those with active CeD and conducted internal transcribed spacer (ITS) sequencing using *Parabasalia* specific primers (*16*). Interestingly, we detected *Parabasalia* in 38% of controls (8/21) but only in 10% of active CeD patients (2/20) **(*P* < 0.05,** Fig. 4D), suggesting that *Parabasalia* are underrepresented in the intestines of persons with CeD compared with healthy controls. We detected *Parabasalia* with ITS sequence homology closest to *Pentatrichomonas hominis* and *Tritrichomonas* sp., including *T. arnold*, *T musculis*, and *T. rainier*. Further stratification of the detected *Parabasalia* strains showed that *Tritrichomonas* sp. was significantly overrepresented in controls compared to CeD patients (Fig. 4E), whereas *Pentatrichomonas hominis* was detected in two control and two CeD patients who also were positive for *Tritrichomonas* sp (Fig. 4F). In contrast to a study that found *Dientamoeba fragilis* in up to 30% of individuals from a healthy cohort in Columbia, South America (*16*), we did not detect *Dientamoeba fragilis* in stool samples collected from patients attending the endoscopic clinic at McMaster University, Canada. It is not well understood how *Parabasalia* colonizes the human gut and whether such colonization is persistent or intermittent. These parameters could contribute to the geographically distinct human gut-colonizing *Parabasalia* strains, hence providing further explanation of how environmental factors contribute to differences in CeD prevalence in otherwise genetically similar populations (*1*). Interestingly, a population-based case-control study found that antibiotic use prior to CeD onset, especially metronidazole, which targets anaerobic microbes and is very potent in eradicating gut protists, was associated with CeD development (*29*). Our findings from these studies with human samples support a role for *Tritrichomonas* sp. colonization, an understudied protist of the human gut microbiome, in protecting against the development of CeD. One promising strategy to promote oral tolerance and prevent or revert LOT in CeD is to use the immunoregulatory potential of commensal gut microbes. Here, we identified an oral-tolerance-promoting protist that can prevent virus-mediated LOT to gluten in a CeD-relevant mouse model (fig. S9B). There is an unmet need to develop effective therapies to prevent LOT to gluten in at-risk individuals and to reverse LOT to gluten in CeD patients. Furthermore, the need for the development of an adjunct treatment to the gluten-free diet is supported by the difficulties in strictly excluding gluten from the diet and the lack of mucosal healing observed in 40% of adults with CeD who maintain a gluten-free diet. Our study will motivate a new line of investigation to use oral tolerance-promoting protists in CeD and potentially other immune-mediated food sensitivities including food allergies.

## Acknowledgments

We thank the *Unified Flow Core* in the Department of Immunology at the University of Pittsburgh for flow cytometry analysis and sorting. We thank the Gnotobiotic Core Facility and the Metabolomics and Lipidomics Core at the University of Pittsburgh for their service. We thank the clinical team at McMaster University. Model figures were created with BioRender.

## Funding

Investigator Start-up Fund, Department of Immunology, University of Pittsburgh School of Medicine (MM, RH)

The Global Grants for Gut Health, co-supported by Yakult and Nature Research (RH)

National Institute of Health grant R21AI163721 (RH)

National Institute of Health grant R01DK098435 (BJ, TSD)

National Institute of Health grant R01DK130897 (MM)

Austrian Marshall Plan Foundation fellowship (MS)

Tsinghua University and Chinese Scholarship Council fellowship (YZ)

Institute for Infection, Inflammation, and Immunity in Children (i4Kids) grant (RH, TSD)

National Institute of Health T32 AI089443 (LVDK)

National Institute of Health T32 AI060525 (PHB)

PACER Catalytic Award (RH)

National Institute of Health grant S10OD023402 (SGW)

Canadian Institutes of Health Research -Project grant 168840 (EVF)

Heinz Endowments (T.S.D.)

## Author contributions

Conceptualization: RH,

Methodology: MS, YZ, LMS, PHB, KAS, MR, JD, PB, HJG, EFV

Investigation: MS, YZ, LMS, PHB, KAS, MR, LVDK, SPP, MJB, BF, LH, SAR, HJG, SJM, SGW, MM, RH

Visualization: KAS, MR, RH

Funding acquisition: SGW, EVF, MM, TSD, BJ, RH

Project administration: RH

Supervision: SCW, PB, JD, EFV, MM, TSD, BJ, RH

Writing original draft: RH

Writing – review and editing: MS, YZ, LMS, MM, TSD, BJ, RH

## Data and materials availability

All data is available in the manuscript or the supplementary materials.

## Materials and Methods

### Human stool samples

Stool samples from consented healthy volunteers (controls) and patients with CeD were collected and processed in anaerobic conditions, frozen, and stored at −80°C until analysis at McMaster University. CeD diagnosis (n=20; age range: 19-75; 60% females) was based on a positive serology for anti-transglutaminase-2 (TG2) antibodies (IgA) or deamidated gliadin antibodies (IgG) and confirmed by duodenal biopsies, as assessed by a pathologist. Controls (n=21; age range:19-57; 55% females) had normal CeD serology and endoscopy and no functional gastrointestinal disorders (according to the Rome IV criteria). Subjects with concomitant inflammatory bowel disease or any other autoimmune disease were excluded. The sample collection was approved by the Hamilton Integrated Research Ethics Board (HiREB #12599-T for CeD patients; HiREB #2820 for controls).

### Mice

All knockout and transgenic mice used in this study are on a C57BL/6 background. RAG^-/-^ OT-II^+/-^ CD45.1^+/+^ mice were provided by Dr. Bana Jabri and bred and housed in our animal facility. HLA-DQ8 transgenic (DQ8tg) mice were described (*10*) and maintained on a gluten-free diet (AIN76A, Envigo). B6.129P2-Il18^tm1Aki^/J (IL-18-deficient) mice and C57BL/6 (WT) mice were purchased from Jackson Laboratories. Female mice were used for experiments. Except for DQ8tg mice which were housed at the University of Chicago, all other mouse strains were housed at the University of Pittsburgh animal facility. In both locations mice were housed under specific pathogen-free (SPF) conditions, where cages were changed on a weekly basis; ventilated cages, bedding, food, and water (non-acidified) were autoclaved before use, ambient temperature maintained at 23°C, and 5% Clidox-S was used as a disinfectant. Experimental cages were randomly housed on two different racks in the vivarium and all cages were kept on automatic 12-h light/dark cycles. Germ-free C57BL/6 WT mice were maintained in flexible film isolators at the University of Pittsburgh Gnotobiotic facility. Animal husbandry for both SPF and germ-free facilities, and experimental procedures were conducted in accordance with Public Health Service policy and approved by the University of Pittsburgh and University of Chicago Institutional Animal Care and Use Committee.

### Infection with reovirus and quantification of virus-specific antibody responses

Reovirus strain type 1 Lang (T1L) virions were purified using CsCl gradient centrifugation and viral titer determinations were conducted using plaque assays as described (*10*). Mice were inoculated perorally with purified 10^9^ plaque-forming units (PFU) of T1L diluted in phosphate-buffered saline (PBS) using a 22-gauge round-tipped needle (Cadence Science). Reovirus-specific antibody responses were determined using a 60% plaque-reduction neutralization assay (PRNT 60) as described (*10*).

### Isolation and colonization with *Tritrichomonas arnold*

Cecal contents were harvested from *T. arnold* positive C57BL/6 mice and mashed through a 100 µM cell strainer. Cecal contents were washed 3 times with PBS containing 0.5 µg/ml Amphotericin, 20 µg/ml Gentamicin, 100 µg/ml Streptomycin, 50 µg/ml Vancomycin and 100 U/ml Penicillin (Sigma) at 1000 rpm for 5 minutes. *T. arnold* were further purified by a 40% Percoll at 1000g for 15 minutes without braking. *T. arnold* were washed 3 times with sterile PBS to remove remaining antibiotics. The number and viability of isolated *T. arnold* was determined by counting with a hemocytometer. Approximately 1×10^6^ *T. arnold* in 200µl PBS were orally gavaged into *Tritrichomonas* negative C57BL/6 mice. For germ-free experiments *T. arnold* was sorted twice on a BD Aria IIU (BD Biosciences) post Percoll purification based on size (forward scatter) to exclude any residual bacterial contaminants.

### *In-vivo* T cell conversion assay

At the start of the experiment C57BL/6 mice were colonized with 1×10^6^ *T. arnold* in 200µl PBS or PBS alone by oral gavage. 12 days post *T. arnold* administration naïve CD4^+^ T cells were purified from the spleen and lymph nodes of RAG^-/-^ OT-II^+/-^ CD45.1^+/+^ mice and sorted on a BD Aria IIU (BD Biosciences). Approximately, 10^5^ OT-II cells were transferred retro-orbitally into congenic C57BL/6 mice. 1 day post OT-II cell transfer, mice were gavaged with 10^9^ PFU T1L in 200µl PBS or PBS alone. Mice were fed an OVA-containing diet (ENVIGO, TD.130362, 10 mg/kg) for 6 days. For Germ-free experiments mice received OVA (grade V, Sigma) dissolved in the drinking water (1.5%) for 6 days.

### *In vivo* succinate treatment

C57BL/6 mice received 150mM succinate in drinking water 12 days prior to *in vivo* T cell conversion assay for the duration of the experiment.

### Loss of tolerance and delayed type hypersensitivity assay

At the start of the experiment a subset of DQ8tg mice was colonized with 1×10^6^ *Tritrichomonas* in 200µl PBS by oral gavage. 12 days post *T. arnold* administration, *mice* received a dose of 50 mg of chymotrypsin-digested gliadin (CT-gliadin) (*10*) by oral gavage for 2 days. In the course of the first CT-gliadin feeding, a subset of mice was gavaged with 10^9^ PFU T1L. Two days after CT-gliadin administration and infection, a mixture of complete Freund’s adjuvant (CFA, Sigma) and CT-gliadin was administered subcutaneously in the lower back as an emulsion of 100 µl CFA and 100 µl PBS containing 300 µg CT-gliadin, under isoflurane gas anesthesia. At day 27, mouse sera were obtained by submandibular bleeding for anti-gliadin IgG2c ELISA quantification. Anti-gliadin IgG2c ELISA was conducted as described (*10*). Ear challenges were conducted 14 and 24 days after immunization. A volume of 20 µl of 100 µg CT-gliadin / PBS was injected under isoflurane gas anesthesia. Ear thickness was measured 2 days after second OVA challenge using a digital precision caliper (Fisher Scientific). Swelling was determined by subtracting pre-challenge from post-challenge ear thickness.

### DNA extraction from ceca for 28S qPCR

Total DNA was extracted from cecal contents using the Fast DNA Stool Mini Kit (Qiagen, 51604). For quantification, quantitative PCR (qPCR) of the 28S rRNA-encoding gene was conducted on a Bio-Rad CFX384 using iTaq^TM^ SYBR (Bio-Rad, 1725125). *Tritrichomonas* 28S rRNA gene qPCR was normalized to the host murine *Ifnb1* gene (*15, 30*).

### Parabasalia internal transcribed spacer (ITS) sequencing

Total DNA was extracted from stool samples using the Fast DNA Stool Mini Kit (Qiagen, 51604). For ITS sequencing of the *Parabasalia* identified in the University of Pittsburgh vivarium and human stool samples the ITS region was PCR-amplified using pan-parabasalid primers described in (*16, 17*). The resulting PCR products were sequenced by Sanger Sequencing (Azenta Life Sciences) and a BLASTn search was conducted. Alignment with the ITS sequences of *T. muris* (GenBank: AY886843.1), *T. musculis* (GenBank: KX000922.1), *T. rainier* (GenBank: MH370486.1), and *Tritrichomonas sp.* (GenBank: MF375342.1) was conducted (*31, 32*).

### RNA processing and RT-PCR

RNA was prepared using the RNeasy Mini Kit (Qiagen, 74136). cDNA synthesis was conducted using iScript^TM^ (Bio-Rad, 1708891BUN) according to the manufacturer’s instructions. Expression analysis was conducted in duplicate via RT– PCR on a Bio-Rad CFX384 using iTaq^TM^ SYBR (Bio-Rad, 1725125). Expression levels were quantified and normalized to *Gapdh* expression. Murine *Gapdh* and *Il27* primers were described (*10*).

### Antibodies and flow cytometry

Single cell suspensions were pelleted and resuspended in FACS buffer (PBS, 2% FBS) for immunostaining and subsequent flow cytometry analysis. Cell suspensions were incubated with Fc Block (BD Biosciences, 553142) before staining with fluorophore-conjugated monoclonal antibodies. All fluorophore-conjugated antibodies used are listed as follows (clone, fluorophore, company, catalog number): CD45.1 (A20, BV480, BD Biosciences, 746666), CD4 (GK1.5, BV650, BD Biosciences, 563232), I-A/I-E (M5/114.15.2, APC-Fire750, BioLegend, 11-5321-82), Foxp3 (FJK-16s, FITC, eBioscience, 11-5773-82), Tbet (4B10, APC, eBioscience, 17-5825-82), Gata3 (TWAJ, PerCPeF710, eBioscience, 46-9966-42), Rorγt (AFKJS-9, PE, eBioscience, 12-6988-82), IFNg (XMG1.2, BV605, BioLegend, 505839), CD11b (M1/70, APCeF780, eBioscience, 47-0112-82), CD45 (30-F11, BV480, BD Biosciences, 566095), CD44 (IM7, APC, BD Biosciences, 559250), CD62L (MEL-14, PE, eBioscience, 12-0621-81), CD45 (30-F11, APC-R700, BD Biosciences, 565478), CD4 (GK1.5, BUV395, BD Biosciences, 563790), CD11b (M1/70, PEcy7, eBioscience, 25-0112-82), CD19 (1D3, BV605, BD Biosciences, 563148), CD3 (17A2, BV605, BD Biosciences, 564009), Ter119 (TER-119, BV605, BD Biosciences, 563323), IL-12p40 (C17.8, PE, eBioscience, 12-7123-82), CD86 (GL-1, AF647, BioLegend, 105020). Zombie NIR Fixable Viability Kit was purchased from BioLegend (423106). Fixable Aqua Dead Cell Stain Kit was purchased from Life Technologies (L34966). CellTrace Violet Cell Proliferation Kit was purchased from Thermo Fisher Scientific (C34557) and used according to manufacturer’s protocol. For analysis of transcription factors and cytokine expression cells were incubated in RPMI media in the presence of 50 ng/ml phorbol 12-myristate 13-acetate (PMA), 500 ng/ml ionomycin (Sigma), 1.3 µl/ml Golgi Stop and 1 µl/ml Golgi Plug (BD Biosciences) for 3 hours at 37°, 5% CO_2_. For intracellular staining cells were permeabilized with the Foxp3 fixation/permeabilization kit from eBioscience (00-5523-00). For IL-12p40 staining, cells were incubated in RPMI media in the presence of 1 µl/ml Golgi Plug for 6 hours at 37°, 5% CO_2_. For intracellular staining cells were permeabilized with the Cytofix/Cytoperm kit from BD Biosciences (554714). Flow cytometry analysis was conducted on a Cytek Aurora (Cytek). Cell sorting was conducted on a BD Aria IIU (BD Biosciences). Data was analyzed with FlowJo (Treestar).

### Cell dissociation and isolation

Small intestinal draining mesenteric lymph nodes (mLN) and Peyer’s Patches were dissected followed by digestion with 1 mg/ml collagenase VIII (Sigma) in a shaking incubator at 37°, 220rpm for 30 minutes. Cells were mashed through a 100 µm cell strainer to obtain a single cell suspension.

### Oral antigen uptake by DCs

OVA was labelled with Alexa Fluor-647 succinimidyl ester according to the manufacturer’s protocol (Molecular Probes). *T, arnold* colonized mice or control mice received 3.2 mg OVA-Alexa Fluor-647 by oral gavage. Mice were euthanized 18 hours post-feeding and OVA uptake by DCs in mLN was assessed by flow cytometry.

### *In vitro* T cell conversion assay

mLN DCs were isolated and purified (CD11c Positive Selection Kit II, STEMCELL Technologies) 2 days after inoculation of WT mice (*T. arnold* colonized or non-colonized) with T1L or PBS. 2×10^4^ DCs were co-cultured for 3 days with 5×10^4^ FACS sorted naïve OT-II CD4+ T cells in the presence of 1 µg/ml OVA peptide (OVA323-339, Invivogen) and 0.25 ng/ml TGFβ (recombinant mouse TGF-b1, BioLegend).

### Analysis of cytokine production

IFNγ was measured using electrochemiluminescence (IFNg V-PLEX Mouse, Meso Scale Diagnostics).

### Measurement of Short Chain Fatty Acids in luminal contents

Cecum samples were homogenized with 50% aqueous acetonitrile at a ratio of 1:15 vol:wt 5 µg/mL Deuterated internal standards: (D_2_)-formate, (D_4_)-acetate, (D_5_)-butyrate, (D_6_)-propionate, (D_2_)-valerate and (D_4_)-hexanoate (CDN Isotopes) were added. Samples were homogenized using a FastPrep-24 system (MP-Bio), with Matrix D at 60hz for 30 seconds, before being cleared of protein by centrifugation at 16,000 x *g*. 60 µL cleared supernatants were collected and derivatized using 3-nitrophenylhydrazine. Each sample was mixed with 20 µL of 200 mM 3-nitrophenylhydrazine in 50% aqueous acetonitrile and 20 µL of 120 mM N-(3-dimethylaminopropyl)-N0-ethylcarbodiimide −6% pyridine solution in 50% aqueous acetonitrile. The mixture was reacted at 50 °C for 40 minutes and the reaction was stopped with 0.45 mL of 50% acetonitrile. Derivatized samples were injected (50 µL) via a Thermo Vanquish UHPLC and separated over a reversed phase Phenomenex Kinetex 150 mm x 2.1 mm 1.7 µM particle C18 maintained at 55°C. For the 20-minute LC gradient, the mobile phase consisted of the following: solvent A (water / 0.1% FA) and solvent B (ACN / 0.1% FA). The gradient was the following: 0-2 minutes 15% B, increase to 60% B over 10 minutes, continue increasing to 100% B over 1 minute, hold at 100% B for 3 minutes, reequillibrate at 15% B for 4 minutes. The Thermo IDX tribrid mass spectrometer was operated in both positive ion mode, scanning in ddMS2 mode (2 μscans) from 75 to 1000 m/z at 120,000 resolution with an AGC target of 2e5 for full scan, 2e4 for ms2 scans using HCD fragmentation at stepped 15,35,50 collision energies. Source ionization setting was 3.0kV spray voltage respectively for positive mode. Source gas parameters were 45 sheath gas, 12 auxiliary gas at 320°C, and 3 sweep gas. Calibration was conducted prior to analysis using the PierceTM FlexMix Ion Calibration Solutions (Thermo Fisher Scientific). Integrated peak areas were then extracted manually using Quan Browser (Thermo Fisher Xcalibur ver. 2.7). SCFA are reported as area ratio of SCFA to the internal standard (*33*).

### Histology for PAS (Periodic acid Schiff staining)

A 10cm piece of the jejunum was removed, cut longitudinally, and pinned down onto wax. Tissue pieces were fixed in 4% paraformaldehyde (PFA, Fischer Scientific) for 3 hours at 4 °C and dehydrated in 30% w/v sucrose overnight. Tissue sections were coiled into a Swiss roll and embedded in optimal cutting temperature (OCT) compound (Tissue-Tek) and stored at −80 °C prior to sectioning. Sections were sliced to 8 µm on a cryostat (Leica CM 1950) and fixed in RT methanol (Sigma-Aldrich) for 2 minutes prior to PAS staining according to manufacturer’s protocol (Sigma-Aldrich, 395B-1KT). Slides were digitized for representative section (Nikon 90i, 0.75 N.A. 20x objective and a Hamamatsu Flash 4.0 CMOS camera) and were quantified for goblet cells per villi or per 100 intestinal epithelial cells.

### Immunofluorescence Staining

For Tuft cell staining, 8 µm Swiss roles were fixed in 4% PFA for 10 minutes at RT. Slides were washed in blocking buffer (1% w/v bovine serum albumin, 0.1% Tween20,1x PBS) for 10 minutes at RT. Sections were surrounded using a hydrophobic barrier (PAP pen, Thermo-Fischer) and permeabilized by incubating tissue in permeabilization buffer (0.4% Tritonx100, 1% fetal bovine serum, 1x PBS) for 10 minutes at RT. Sections were then incubated with anti-mouse IgG DCAMKL1 antibody (Abcam, ab31704) in blocking buffer for 1 hour at RT, washed three times in PBS, and incubated with anti-rabbit IgG (H+L) antibody, Alexa Flour Plus 647 (Thermo Fisher Scientific, A32733) for 30 minutes at RT, and washed 3 times in PBS. Slides were incubated in blocking buffer for 30 minutes at RT prior to staining for epithelial cells by anti-mouse IgG2a CD326/Ep-CAM conjugated to Alexa-Fluor 488 (Biolegend, 118210) for 1 hour at RT, washed three times in PBS, counter stained with 500 ng/ml 4’,6-Diamidino-2-Phenylindole, Dihydrochloride (DAPI, Thermofisher) for 5 minutes at RT, washed three times in PBS, and mounted in Fluoromount-G (Thermofisher). Slides were stored light protected at 4 °C until digitized (Nikon Ti inverted microscope equipped with a Nikon A1 point scanning confocal scan head and a 1.40 N.A. 20x Objective). Tuft cells were quantified for tuft cells per villi.

### Visualization and quantification of TG2 activity

At the start of the experiment DQ8tg mice were colonized with 10^6^ *Tritrichomonas* in 200µl PBS or PBS alone by oral gavage. 12 days post *T. arnold* administration mice were gavaged with 10^9^ PFU T1L in 200µl PBS or PBS alone. The *in vivo* TG2 enzymatic activity was assessed 18 hours post T1L infection. Six and three hours prior to euthanasia, mice were injected i.p. (100 mg/kg) with 5-(biotinamido)-pentylamine (5-BP), a substrate for TG2 transamidation activity, which was synthesized following a published protocol (*34*). Jejunal pieces were collected and frozen in optimum cutting temperature (OCT) compound (Tissue-Tek). Frozen sections of 5 µm thickness were cut, fixed in 1% paraformaldehyde, and TG2 protein was visualized by staining with a rabbit polyclonal anti-TG2 antibody (custom produced by Pacific Immunology), followed by AF488-conjugated donkey ant-rabbit IgG antibody (Invitrogen, A32790). TG2 enzymatic activity was measured using 5-BP crosslinking and was visualized by costaining with AF594-conjugated streptavidin (Invitrogen, S32356). Images were acquired at 20x magnification using a Leica SP8 Laser Scanning Confocal microscope. TG2 activity was quantified by systematically taking two sections of the jejunum from each mouse, quantifying the 5-BP signal / TG2 protein signal on a per villi basis. The mean 5-BP signal / TG2 signal is shown for each mouse that was assessed.

### RNA sequencing processing and data analysis

RNA sequencing libraries were prepared using the Illumina TruSeq protocol and sequenced with paired-end 75 bp reads on an Illumina HiSeq. FASTQ sequences with Phred scores less than 20 were trimmed from 3 prime end and reads smaller than 25 bp were removed. The remaining clean reads were aligned to the mm10 protein-coding reference library from Ensembl using STAR. Noise reduction filter was applied to quantified transcript counts to exclude features with maximum counts less than 10. DEseq2 was used to normalize the read count data and to assess for differential expression. Genes with absolute value of |log2(fold change)| >2 and false discovery rate <0.1 were considered as significant differentially expressed genes (DEGs).

### Statistical analysis

Mice were allocated to experimental groups based on their genotype and randomized within the given age-matched groups. Male and female mice were used. Because our mice were inbred and matched for age and sex, we always assumed similar variance between the different experimental groups. All experimental and control animals were littermates, and none were excluded from the analysis at the time of collection. Data were analyzed using unpaired two-tailed Student’s *t*-test for single comparisons, and one-way ANOVA analysis for multiple comparisons. ANOVA analysis was followed by a Tukey’s post hoc test. Chi-squared test was used test significance of observed frequencies of *Parabasalia* in human stool samples. Figures and statistical analysis were generated using GraphPad Prism 8 (GraphPad Software). The statistical test used, and *P* values are indicated in each figure legend. *P* values of < 0.05 were considered statistically significant. \**P* < 0.05, \*\**P* < 0.01, \*\*\**P* < 0.001 and \*\*\*\**P* < 0.0001. ns = not significant.

---

**Fig. S1.**
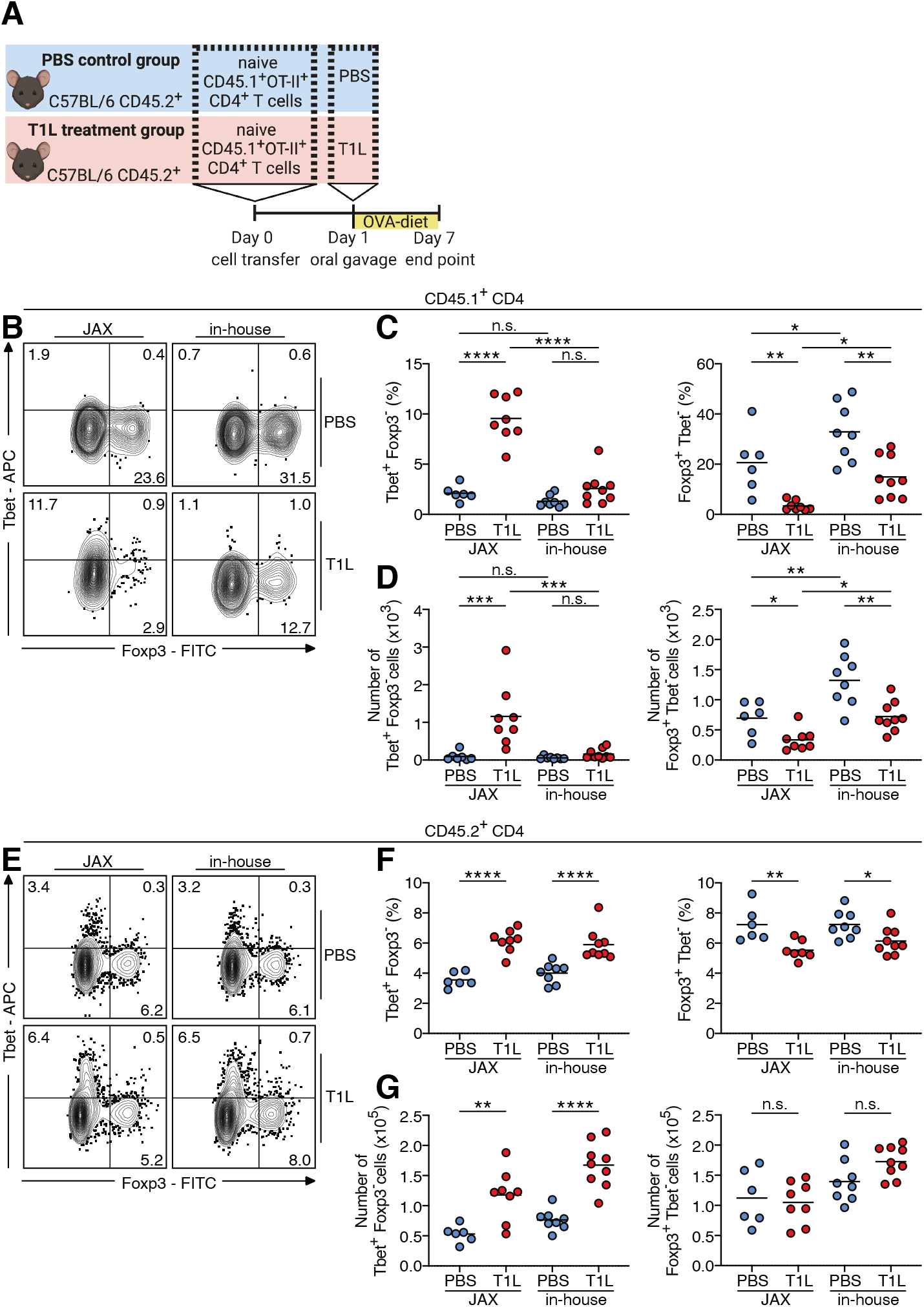
Suppression of T1L-induced loss of oral tolerance is associated with the presence of *Tritrichomonas* in in-housed mice. **(A)** Schematic for OT-II T cell conversion experiments described in Fig. 1A-C. **(B-G)** Naïve OT-II^+^ CD45.1^+^ CD4^+^ T cells were transferred into WT CD45.2^+^ mice. One day after transfer mice were inoculated perorally with 10^9^ PFU of T1L or PBS and fed an OVA-containing diet for 6 days. The intracellular expression of Tbet and Foxp3 in (B-D) transferred OT-II^+^ CD45.1^+^ CD4^+^ T cells (E-G) host CD45.2^+^ CD4^+^ T cells in the mLN was evaluated by means of flow cytometry. (B and E) Representative contour plots, (C and F) percentages, and (D and G) absolute numbers are shown. Center is mean, one-way ANOVA, Tukey’s post hoc test. \**P* < 0.05, \*\**P* < 0.01, \*\*\**P* < 0.001, \*\*\*\**P* < 0.0001, n.s. not significant.

**Fig. S2.**
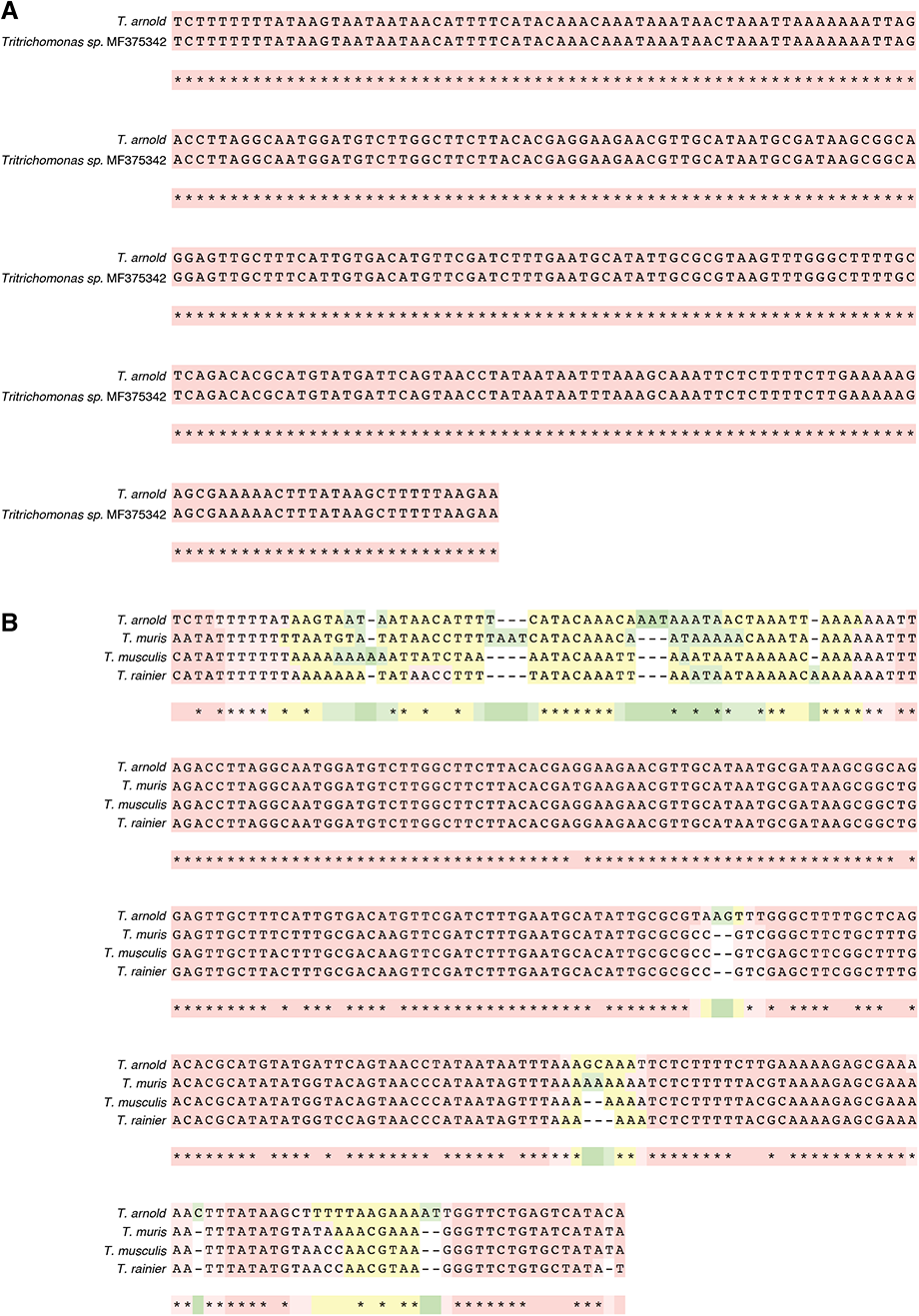
Alignment of *Tritrichomonas* strains. Alignment of ITS of *T. arnold* with published sequences for **(A)** *Tritrichomonas sp.* MF375342 and **(B)** *T. muris*, *T musculis*, and *T. rainier*.

**Fig. S3.**
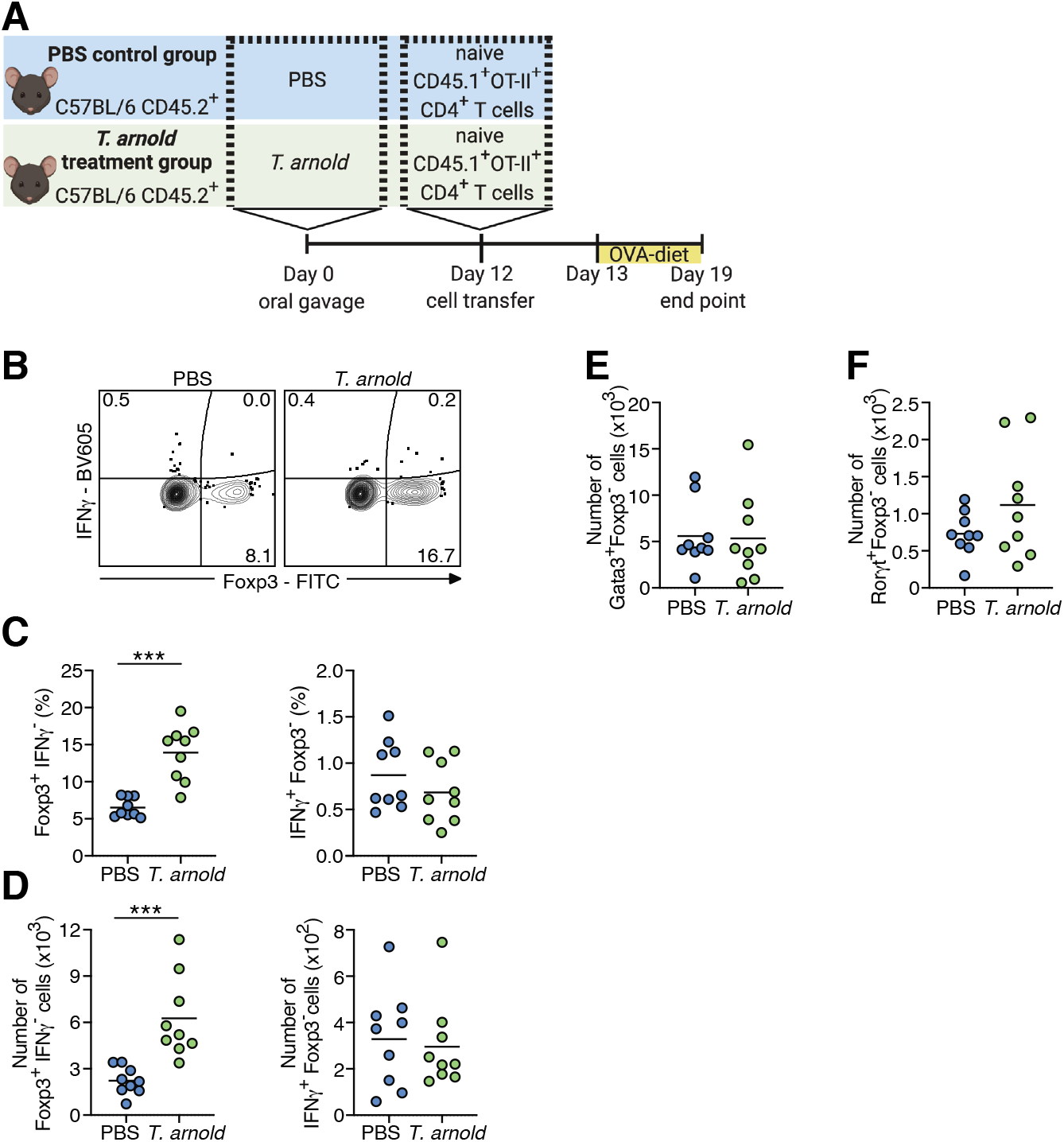
*T. arnold* promotes dietary-antigen specific-Treg responses. **(A)** Schematic for OT-II T cell conversion experiments described in Supplementary Figure S3B-F. **(B-F)** WT CD45.2^+^ mice were inoculated perorally with 10^6^ *T. arnold* or PBS for 12 days prior to transfer of naïve OT-II^+^ CD45.1^+^ CD4^+^ T cells into WT CD45.2^+^ mice. One day after transfer mice were fed an OVA-containing diet for 6 days. The intracellular expression of (B-D) Foxp3 and IFNγ, (E) Gata3, and (F) Rorγt in transferred OT-II^+^ CD45.1^+^ CD4^+^ T cells in the mLN was evaluated by means of flow cytometry. (B) Representative contour plots, (C) percentages and (D-F), absolute numbers are shown. (C-F) Data are representative of two independent experiments (n = 9 both groups). Center is mean, two-tailed unpaired *t*-test. \*\*\**P* < 0.001.

**Fig. S4.**
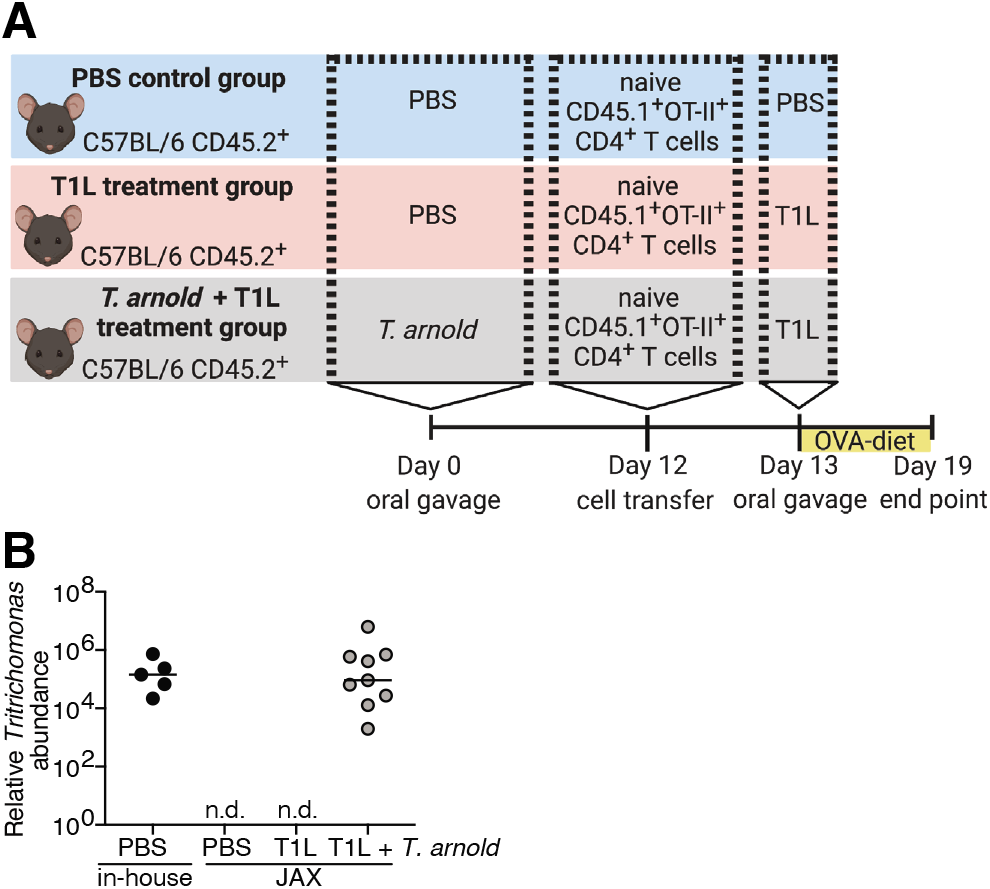
*T. arnold* protects against T1L-mediated loss of oral tolerance. **(A)** Schematic for OT-II T cell conversion experiments described in Fig. 2A-C. **(B)** WT CD45.2^+^ mice were inoculated perorally with 10^6^ *T. arnold* or PBS for 12 days followed by peroral inoculation with 10^9^ PFU of T1L or PBS for 6 days. *T. arnold* gavaged and naturally colonized in-housed mice were quantified by qPCR using DNA isolated from cecal contents. 28S normalized to the host murine *Ifnb1* gene. Median is shown.

**Fig. S5.**
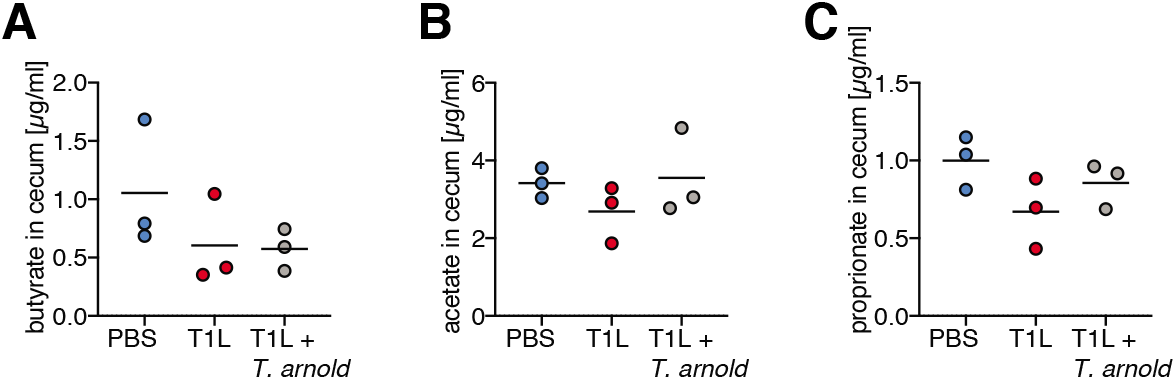
*T. arnold* colonization does not alter SCFA abundance in the intestine. **(A and C)** WT mice were inoculated perorally with 10^6^ *T. arnold* or PBS for 12 days prior to T1L infection*. T. arnold* colonized mice or control mice were inoculated perorally with 10^9^ PFU of T1L or PBS for 6 days. Levels of (A) butyrate, (B) acetate, and (C) propionate were measured by LC-MS in cecal contents of T1L infected mice in the presence or absence of *T. arnold* colonization or controls and are displayed as µg per ml (n = 3 mice per group). Center is mean.

**Fig. S6.**
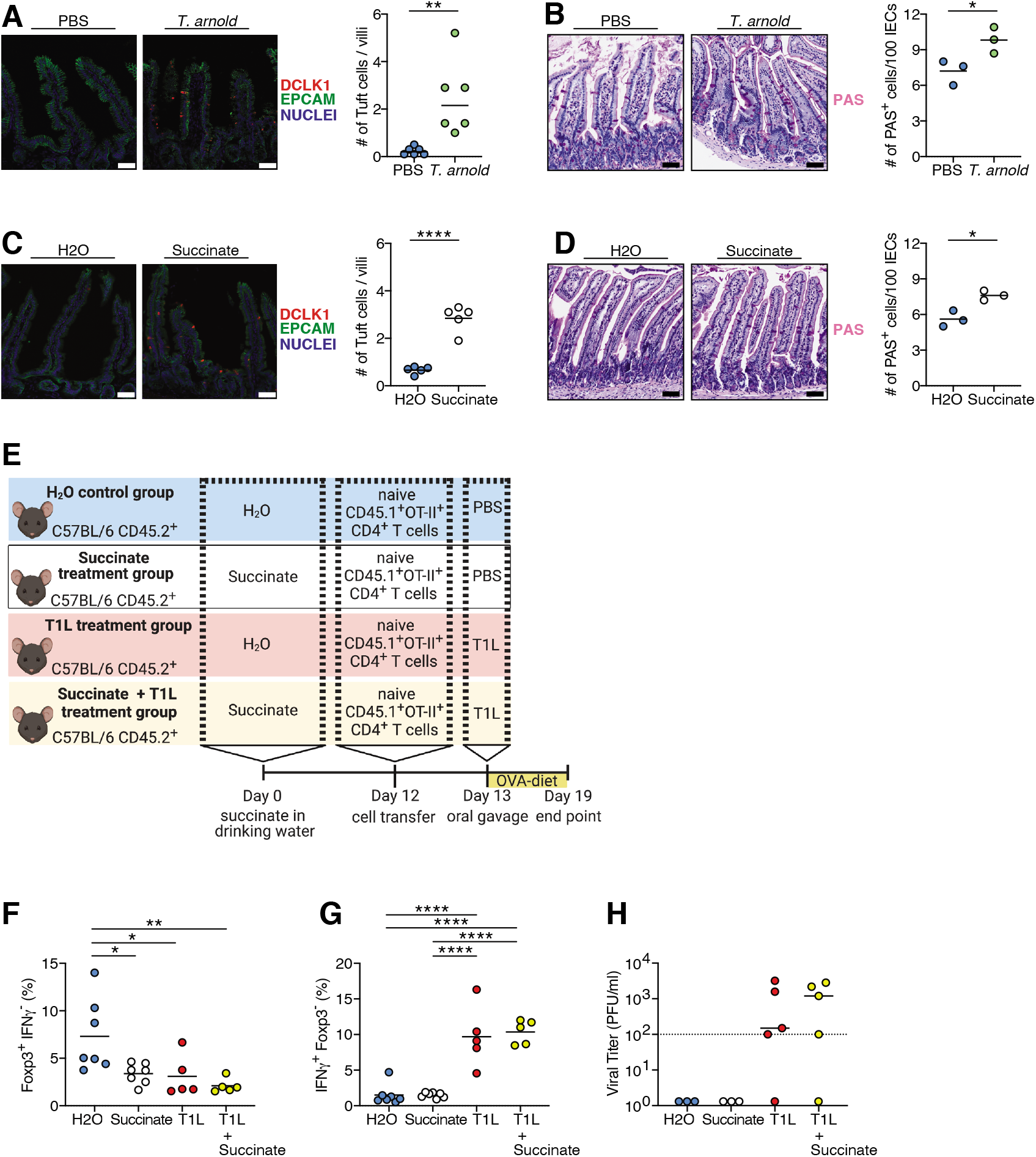
Succinate is not sufficient to protect against T1L-mediated inflammatory responses to dietary antigens. **(A and B)** WT mice were inoculated perorally with 10^6^ *T. arnold* or PBS for 12 days. (A) Presence of tuft cells in the jejunum visualized by the expression of DCLK1 in red; epithelial cells (EPCAM) in green; DAPI in blue. (B) Total number of PAS^+^ goblet cells per 100 intestinal epithelial cells (IECs) in the jejunum was determined. (A-B) Representative images (left panel), quantification (right panel); Scale bars, 50 µm. **(C-D)** WT mice received 150mM succinate in drinking water or regular drinking water for 12 days. (C) Presence of tuft cells in the jejunum visualized by the expression of DCLK1 in red; epithelial cells (EPCAM) in green; DAPI in blue. (D) Total number of PAS^+^ goblet cells per 100 IECs in the jejunum was determined. (C-D) Representative images (left panel), quantification (right panel); Scale bars, 50 µm. **(E-H)** WT mice received 150mM succinate in drinking water or regular drinking water for 12 days followed by an OT-II T cell conversion assay. Naïve OT-II^+^ CD45.1^+^ CD4^+^ T cells were transferred into WT CD45.2^+^ mice. One day after transfer mice were inoculated perorally with 10^9^ PFU of T1L or PBS and fed an OVA-containing diet for 6 days. (F-G) The intracellular expression of Foxp3 and IFNγ in transferred OT-II^+^ CD45.1^+^ CD4^+^ T cells in the mLN was evaluated by means of flow cytometry. Percentages are shown. **(H)** At 6 days post infection, a 3cm section of ileum was resected, and viral titers in this tissue were determined by plaque assay. (A-D) Center is mean, two-tailed unpaired *t*-test (n = 3-6 mice per group). (F-H) Data are representative of two independent experiments (n = 5-7 mice per group). (F-G) Center is mean, one-way ANOVA, Tukey’s post hoc test. (H) Center is median. \**P* < 0.05, \*\**P* < 0.01, \*\*\**P* < 0.01, \*\*\*\**P* < 0.0001.

**Fig. S7.**
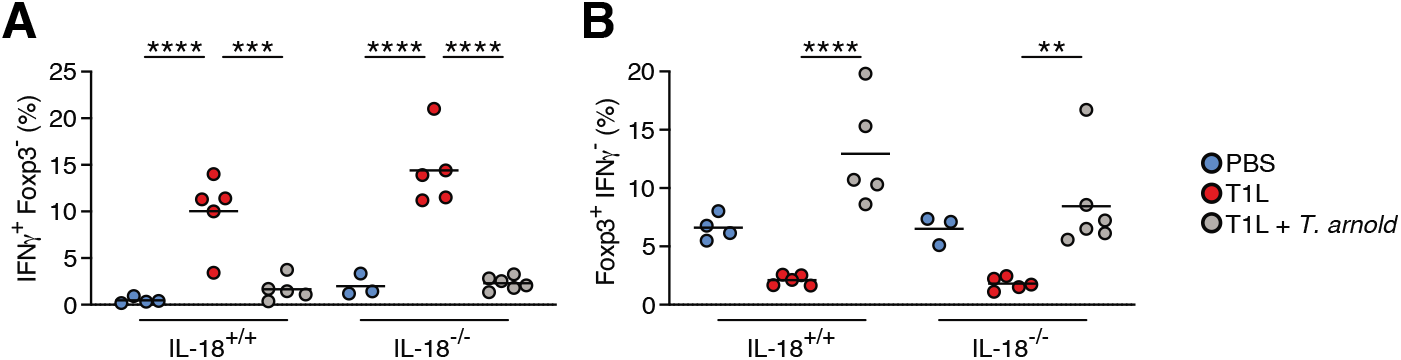
IL-18 is dispensable for *T. arnold*-mediated protection against T1L-mediated inflammatory responses to dietary antigens. **(A and B)** IL-18^-/-^ CD45.2^+^ mice and WT controls were inoculated perorally with 10^6^ *T. arnold* or PBS for 12 days prior to OT-II T cell transfer. One day after transfer mice were inoculated perorally with 10^9^ PFU of T1L or PBS and fed an OVA-containing diet for 6 days. The intracellular expression of Foxp3 and IFNγ in transferred OT-II^+^ CD45.1^+^ CD4^+^ T cells in the mLN was evaluated by means of flow cytometry. Percentages are shown. (A-B) Center is mean, one-way ANOVA, Tukey’s post hoc test. \*\**P* < 0.01, \*\*\**P* < 0.01, \*\*\*\**P* < 0.0001.

**Fig. S8.**
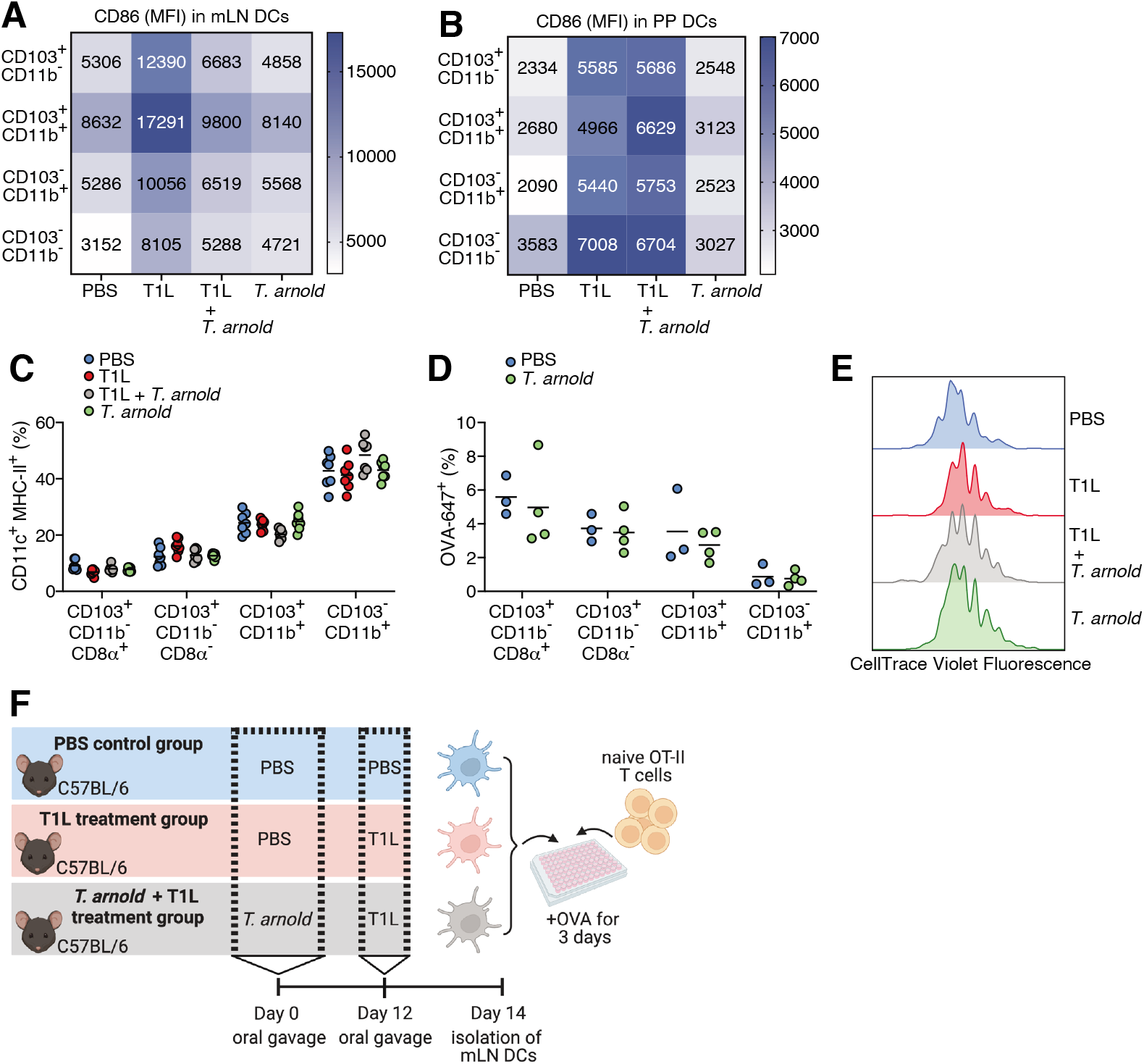
*T. arnold* suppresses T1L-induced DC activation in mLN but not PP. **(A-B)** WT mice were inoculated perorally with 10^6^ *T. arnold* or PBS for 12 days followed by an oral infection with 10^9^ PFU T1L or PBS for 48 hours. The expression of CD86 (A) in mLN DC subsets and (B) PP DC subsets were evaluated by flow cytometry. Mean fluorescence intensity (MFI) is displayed in heatmap. Mean MFI of n = 3-7 per is shown. **(C-D)** Mice were colonized with 10^6^ *T. arnold* for 12 days or PBS control. (C) Frequencies of mLN DC subsets were assessed by means of flow cytometry. (D) Mice were gavaged with OVA-Alex Fluor-647 18 hours before euthanasia and OVA-Alex Fluor-647 uptake by DCs in the mLN was analyzed by flow cytometry. Percentages of OVA-Alex Fluor-647 uptake are shown in indicated DC subsets. Center is mean. **(E)** WT CD45.2^+^ mice were inoculated perorally with 10^6^ *T. arnold* or PBS for 12 days prior to transfer of CellTrace Violet labelled naïve OT-II^+^ CD45.1^+^ CD4^+^ T cells into WT CD45.2^+^ mice. One day after transfer mice were inoculated perorally with 10^9^ PFU of T1L or PBS and fed an OVA-containing diet for 3 days. CellTrace Violet dye dilution was measured as a proxy for proliferation. Representative histograms from n = 4 per treatment group are shown. **(F)** Schematic of mLN DC-T cell co-culture experiments described in Fig. 3H-I. (A-E) Data are representative of two independent experiments (individual dots represent n-numbers).

**Fig. S9.**
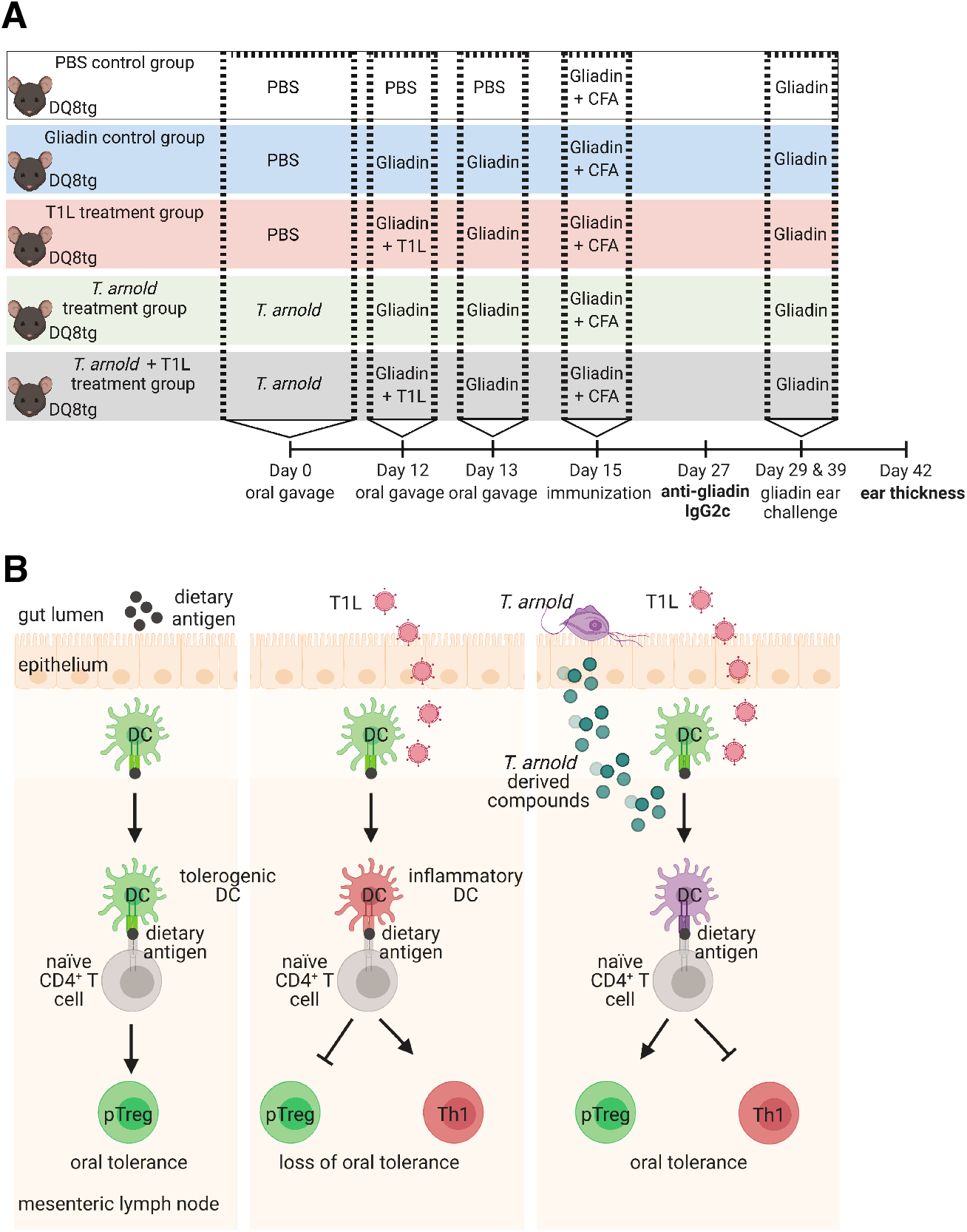
*Parabasalia* protect against loss of oral tolerance to gluten. **(A)** Schematic for delayed type hypersensitivity (DTH) experiments described in Fig. 4A-B. **(B)** Under homeostatic conditions, regulatory T cell (pTreg) responses to dietary antigen are induced in the mesenteric lymph nodes draining the small intestine (oral tolerance). T1L reovirus infection promotes inflammatory responses in dietary-antigen-presenting dendritic cells (DC). Consequently, T1L infection inhibits the conversion of pTregs and promotes development of T helper 1 (Th1) immunity to dietary antigen (loss of oral tolerance). Colonization with the protist *Tritrichomonas (T.) arnold* prevents T1L-mediated inflammatory responses in dietary-antigen-presenting DCs. Consequently, *T. arnold* prevents T1L-mediated loss of oral tolerance by restoring pTreg responses and preventing Th1 immunity to dietary antigens.

## References and Notes

1. V. Abadie, L. M. Sollid, L. B. Barreiro, B. Jabri, Integration of genetic and immunological insights into a model of celiac disease pathogenesis. Annu Rev Immunol 29, 493–525 (2011).

2. J. M. Tjon, J. van Bergen, F. Koning, Celiac disease: how complicated can it get? Immunogenetics 62, 641–651 (2010).

3. L. M. Sollid, B. Jabri, Triggers and drivers of autoimmunity: lessons from coeliac disease. Nat Rev Immunol 13, 294–302 (2013).

4. E. M. Nilsen et al., Gluten induces an intestinal cytokine response strongly dominated by interferon gamma in patients with celiac disease. Gastroenterology 115, 551–563 (1998).

5. B. Jabri, L. M. Sollid, Tissue-mediated control of immunopathology in coeliac disease. Nat Rev Immunol 9, 858–870 (2009).

6. L. Plot, H. Amital, Infectious associations of Celiac disease. Autoimmun Rev 8, 316–319 (2009).

7. S. L. Smits et al., Human astrovirus infection in a patient with new-onset celiac disease. J Clin Microbiol 48, 3416–3418 (2010).

8. L. C. Stene et al., Rotavirus infection frequency and risk of celiac disease autoimmunity in early childhood: a longitudinal study. Am J Gastroenterol 101, 2333–2340 (2006).

9. R. Troncone, S. Auricchio, Rotavirus and celiac disease: clues to the pathogenesis and perspectives on prevention. J Pediatr Gastroenterol Nutr 44, 527–528 (2007).

10. R. Bouziat et al., Reovirus infection triggers inflammatory responses to dietary antigens and development of celiac disease. Science 356, 44–50 (2017).

11. O. Pabst, A. M. Mowat, Oral tolerance to food protein. Mucosal Immunol 5, 232–239 (2012).

12. D. Esterhazy et al., Compartmentalized gut lymph node drainage dictates adaptive immune responses. Nature 569, 126–130 (2019).

13. J. von Moltke, M. Ji, H. E. Liang, R. M. Locksley, Tuft-cell-derived IL-25 regulates an intestinal ILC2-epithelial response circuit. Nature 529, 221–225 (2016).

14. F. Gerbe et al., Intestinal epithelial tuft cells initiate type 2 mucosal immunity to helminth parasites. Nature 529, 226–230 (2016).

15. M. R. Howitt et al., Tuft cells, taste-chemosensory cells, orchestrate parasite type 2 immunity in the gut. Science 351, 1329–1333 (2016).

16. A. Chudnovskiy et al., Host-Protozoan Interactions Protect from Mucosal Infections through Activation of the Inflammasome. Cell 167, 444–456 e414 (2016).

17. M. S. Nadjsombati et al., Detection of Succinate by Intestinal Tuft Cells Triggers a Type 2 Innate Immune Circuit. Immunity 49, 33–41 e37 (2018).

18. N. Arpaia et al., Metabolites produced by commensal bacteria promote peripheral regulatory T-cell generation. Nature 504, 451–455 (2013).

19. C. Schneider et al., A Metabolite-Triggered Tuft Cell-ILC2 Circuit Drives Small Intestinal Remodeling. Cell 174, 271–284 e214 (2018).

20. W. Lei et al., Activation of intestinal tuft cell-expressed Sucnr1 triggers type 2 immunity in the mouse small intestine. Proc Natl Acad Sci U S A 115, 5552–5557 (2018).

21. J. R. McDole et al., Goblet cells deliver luminal antigen to CD103+ dendritic cells in the small intestine. Nature 483, 345–349 (2012).

22. D. H. Kulkarni et al., Goblet cell associated antigen passages support the induction and maintenance of oral tolerance. Mucosal Immunol, (2019).

23. M. N. Fleeton et al., Peyer’s patch dendritic cells process viral antigen from apoptotic epithelial cells in the intestine of reovirus-infected mice. J Exp Med 200, 235–245 (2004).

24. D. Esterhazy et al., Classical dendritic cells are required for dietary antigen-mediated induction of peripheral T(reg) cells and tolerance. Nat Immunol 17, 545–555 (2016).

25. R. Hinterleitner, B. Jabri, A dendritic cell subset designed for oral tolerance. Nat Immunol 17, 474–476 (2016).

26. J. L. Coombes et al., A functionally specialized population of mucosal CD103+ DCs induces Foxp3+ regulatory T cells via a TGF-beta and retinoic acid-dependent mechanism. J Exp Med 204, 1757–1764 (2007).

27. K. M. Luda et al., IRF8 Transcription-Factor-Dependent Classical Dendritic Cells Are Essential for Intestinal T Cell Homeostasis. Immunity 44, 860–874 (2016).

28. M. Martinez-Lopez, S. Iborra, R. Conde-Garrosa, D. Sancho, Batf3-dependent CD103+ dendritic cells are major producers of IL-12 that drive local Th1 immunity against Leishmania major infection in mice. Eur J Immunol 45, 119–129 (2015).

29. K. Marild et al., Antibiotic exposure and the development of coeliac disease: a nationwide case-control study. BMC Gastroenterol 13, 109 (2013).

30. M. Meisel et al., Microbial signals drive pre-leukaemic myeloproliferation in a Tet2-deficient host. Nature 557, 580–584 (2018).

31. P. Di Tommaso et al., T-Coffee: a web server for the multiple sequence alignment of protein and RNA sequences using structural information and homology extension. Nucleic Acids Res 39, W13–17 (2011).

32. C. Notredame, D. G. Higgins, J. Heringa, T-Coffee: A novel method for fast and accurate multiple sequence alignment. J Mol Biol 302, 205–217 (2000).

33. J. Han, K. Lin, C. Sequeira, C. H. Borchers, An isotope-labeled chemical derivatization method for the quantitation of short-chain fatty acids in human feces by liquid chromatography-tandem mass spectrometry. Anal Chim Acta 854, 86–94 (2015).

34. T. R. DiRaimondo et al., Elevated transglutaminase 2 activity is associated with hypoxia-induced experimental pulmonary hypertension in mice. ACS Chem Biol 9, 266–275 (2014).

